# Infant Sleep Quality and its’ Several Related Factors: An Indonesian Samples

**DOI:** 10.1101/618058

**Authors:** Fiona Muskananfola, Tjhin Wiguna, Raden Irawati Ismail, Teresia Putri Widia Nugraheni, Shafira Chairunnisa

## Abstract

**Objective:** This research aimed to identify factors influencing infant sleep quality including mother-infant bonding, infant stress, parity, and maternal depression and anxiety, and to analyze possible associations between these variables.

**Method:** A cross-sectional design was adopted to analyze data from mothers and their infants (0–36 months of age) using consecutive sampling. Mothers completed two of several questionnaires in the Indonesian language, including the Mother-Infant Bonding Scale, the Brief Infant Sleep Questionnaire, the Symptoms Checklist-90, and an infant stress questionnaire that was specifically designed for this study. The chi-squared test for bivariate analysis and logistic regression were applied to obtain odds ratios for the predictor variables using SPSS version 21 (IBM Corporation, Armonk, NY, USA) for Mac (Apple Inc, Cupertino, CA, USA).

**Results:** Findings indicated that the proportion of infants with problematic sleep quality was 33.30%. Three predictors were significantly associated with problematic infant sleep quality: mother-infant bonding; infant stress; and parity. However, logistic regression analysis demonstrated that only mother-infant bonding (odds ratio [OR] 1.66 [95% confidence interval (CI) 1.15–6.12]) and infant stress (OR 1.29 [95% CI 1.07–2.68]) predicted a 38.7% risk for problematic infant sleep quality.

**Conclusion:** Results of the present study indicated that early detection of mother-infant bonding levels and infant stress is very important. It may be valuable to screen sleep habits for better prognosis among infants because good sleep quality is crucial for optimal growth and development. Results of this study will raise awareness of the importance of mother-infant bonding, infant stress, and problematic infant sleep quality.

## Introduction

Sleep is a reversible state of unresponsiveness from one’s surroundings and is essential for children’s health, development, and daily functions [1]. Lower quality sleep can be indicated by total sleep duration < 9 h, > 3 night awakenings, or nocturnal wakefulness > 1 h [1,2]. Sleep is categorized into two phases based on polysomnography patterns represented by rapid eye movement and non-rapid eye movement. There are 2 processes required in sleep: the homeostatic process (process S); and the circadian rhythm (process C). Process S is required to control the length and depth of sleep, while process C is required for the regulation of sleep in internal organization, and the timing and duration of the sleep-wake cycle. Process S requires the accumulation of somnogens, such as adenosine. Process C acts on environmental cues to synchronize with the 24 h daily cycle [3].

Sleep patterns change as children age. Neonates generally sleep 16–18 h distributed throughout the day. By 1 year of age, the mean duration of night sleep is 10–11 h, and the mean duration of sleep during the day is 2–3 h. The number of sleep hours decreases until 2 years of age and starts to become more concentrated through the night. Hereafter, infant sleep patterns start to become more like that of an adult [1]. The transactional theory of sleep posits that internal and external factors may affect an infant’s sleep quality. Factors include the infant’s health, infant-caretaker relations, culture, environment, and family [4]. Depressive symptoms in mothers demonstrate a prospective relationship with more frequent infant night awakenings [5]. An internal infant factor is stress level, which may impact sleep quality through the cortisol (CT) awakening response (CAR) [6]. Circadian rhythm changes are known to occur during the first 2–3 months of life and regulated similar to those of adults [7]. Sex-based sleep quality differences in previous studies have revealed that females were more sensitive to sleep hormone(s) [8]. External factors, such as parity, is suggested to correlate with infant sleep quality, resulting from the mother’s experiences in parenting [9].

Although sleep is known to be important for infants and toddlers, few studies have investigated infant sleep and the factors that affect it, especially in Asia. Thus, this research aimed to elaborate the probability of several risk factors, such as mother-infant bonding, infant stress, and maternal depression and anxiety in infant sleep quality. The results are anticipated to contribute to the development of methods to improve infant sleep quality.

## Materials and Methods

### Study design and population

This was a cross-sectional study including 63 mother-child dyads with infants (0–36 months of age). The mothers’ passed junior high school, spoke the Indonesian language, and were residents of Jakarta and Depok, Indonesia. Research subjects were recruited using the consecutive sampling technique. Subjects were recruited at the RSCM Kiara Poli Pediatri Sosial, Kampung Lio, and Pondok Labu. An informed consent was obtained from all the parents after the research purpose, procedures, potential risk, the right to withdraw from the study, and confidentiality requirements were fully disclosed by a member of the research team. The research protocol was approved by the Health Research Ethics Committee of Medicine Faculty at Universitas Indonesia (Jakarta, Indonesia).

### Measurements

There were 4 questionnaires used in this study: a basic demographic questionnaire; the Mother-Infant Bonding Scale, Indonesian version I (MIBS-I); the Brief Infant Sleep Questionnaire (BISQ); and the Symptoms Checklist-90 (SCL-90). Mother-infant bonding was measured using the MIBS-I, which was translated and validated in the Indonesian language, and was modified to use among toddlers (0–36 months of age) [10]. It consists of 10 questions using the Likert scoring system, in which “not at all” = 0 and “very much so”/”most of the time” = 3 (Cronbach’s alpha = 0.535). The only items used were from 3 through 10 [10]. Mother-infant bonding was considered abnormal if the MIBS-I Z-score was > 50th percentile.

Infant sleep quality was assessed using the Indonesian version of the BISQ, which has been validated using actigraphy, and is considered to be a reliable tool for sleep problem assessment and screening. The questionnaire consists of sleep ecology, the measurement of sleep, and the parents’ perception of their toddler’s sleep and sleep problems. Mothers were asked to recall the toddlers’ sleep pattern over the past 2 weeks. Infant sleep quality was considered to be disturbed if the toddler slept < 9 h, experienced > 3 night awakenings, or > 1 h of nocturnal wakefulness [2].

Maternal depressive and anxiety symptoms were assessed using the SCL-90, a self-administered questionnaire (Cronbach’s alpha, 0.98), which is consists of 90 statements. Mothers were asked to recall any symptoms of sleep disturbance in the past month. Each statement was scored using a five-point Likert scoring system (i.e., 0–4), in which 0 indicated “not at all” and 4 indicated “often”. The higher the score, the more likely the respondent was to have psychopathological symptoms.

Infant stress level was determined by interviewing the mother according to stress symptoms in the toddler questionnaire such as alteration of sleeping pattern, alteration of eating pattern, emotional symptoms, behavioral symptoms, and measuring the toddler’s heart rate and respiratory rate. This questionnaire consists of 6 questions with ‘yes’ or ‘no’ answers, such as alteration of sleeping pattern (<10 h or >19 h), alteration of eating patterns, emotional symptoms (frequent crying, tantrums, anger, aggressiveness), behavioral symptoms (frequent nail and hair biting), heart rate elevation (> 160 beats/min), and respiratory rate elevation (> 60 breaths/min) [11]. Z-scores from the total number of the above questions that were > 50th percentile were defined as abnormal stress levels.

In addition, the study also collected demographic information from the dyads including age, educational background and parity, among others, using a specific questionnaire that was designed specifically for this study.

### Statistical analysis

Data were stored and analyzed using SPSS version 23 (IBM Corporation, Armonk, NY, USA). Each variable was grouped into 2 categories. The chi-square test was used for bivariate analysis and logistic regression was used to elaborate predictors of problematic infant sleep quality.

## Results

In total, 63 mothers (18–43 years of age) of toddlers 1–36 months of age (55.6% male) completed the questionnaires in full. Mothers were multipara (54.0%), housewives (84.1%), senior high-school graduates (60.3%) who lived in a middle socioeconomic class (42.9%) (Table 1). Toddlers’ sleep conditions and characteristics reported by the parents included: slept in parents’ room (either room sharing or bed sharing, 87.7%); combined sleep position (52.4%); and were helped by feeding in the sleep process (61.9%). More than one-half (54%) of the toddlers did not snore during sleep and woke up at night due to hunger (60.3%) (Table 3). In total, 65% of parents considered their toddler’s sleep as good or excellent (46% and 19%, respectively) (Table 2 and Table 3). However, 42.9% of parents, did not consider the “sleep problem” to be an actual problem. Based on sleep disturbance criteria, 33.3% of the infants in this study were considered to have problematic sleep quality.

**Table 1.** Characteristics of the research subjects (n=63)

| Characteristic | n (%) | Mean ± SD | Median<br>(Minimum-Maximum) |
| --- | --- | --- | --- |
| Infant age, months | | 19.32±10.05 | |
| Infant sex |  |  |  |
| Male | 35(55.6) |  |  |
| Female | 28(44.4) |  |  |
| Infant weight and height |  |  |  |
| Birth weight (g) |  |  | 3.000<br>(1.050-4.900) |
| Mean weight (kg) | | 9.69± 2.93 | |
| Birth length (cm) |  |  | 49.00<br>(33.00-66.00) |
| Mean height (cm) | | 78.59±13.22 | |
| Mother's age (years) |  |  | 29.00<br>(18.00-43.00) |
| Mother education |  |  |  |
| Junior high | 8 (12.7) |  |  |
| Senior high | 38 (60.3) |  |  |
| Bachelor degree | 14 (22.2) |  |  |
| Postgraduate | 3 (4.8) |  |  |
| Mother's occupation |  |  |  |
| Civil servant | 1 (1.6) |  |  |
| Private employee | 5 (7.9) |  |  |
| Professional (teacher/<br>lecturer/farmer/<br>doctor/army officer) | 4 (6.3) |  |  |
| Housewife | 53 (84.1) |  |  |
| Parity |  |  |  |
| Primipara | 29 (46.0) |  |  |
| Multipara | 34 (54.0) |  |  |
| Delivery |  |  |  |
| Spontaneous | 35 (55.6) |  |  |
| C-section | 28 (44.4) |  |  |
| Socioeconomic background |  |  |  |
| Poor | 3 (4.8) |  |  |
| Lower | 8 (12.7) |  |  |
| Middle | 27 (42.9) |  |  |
| Upper middle | 13 (20.6) |  |  |

**Table 2.**
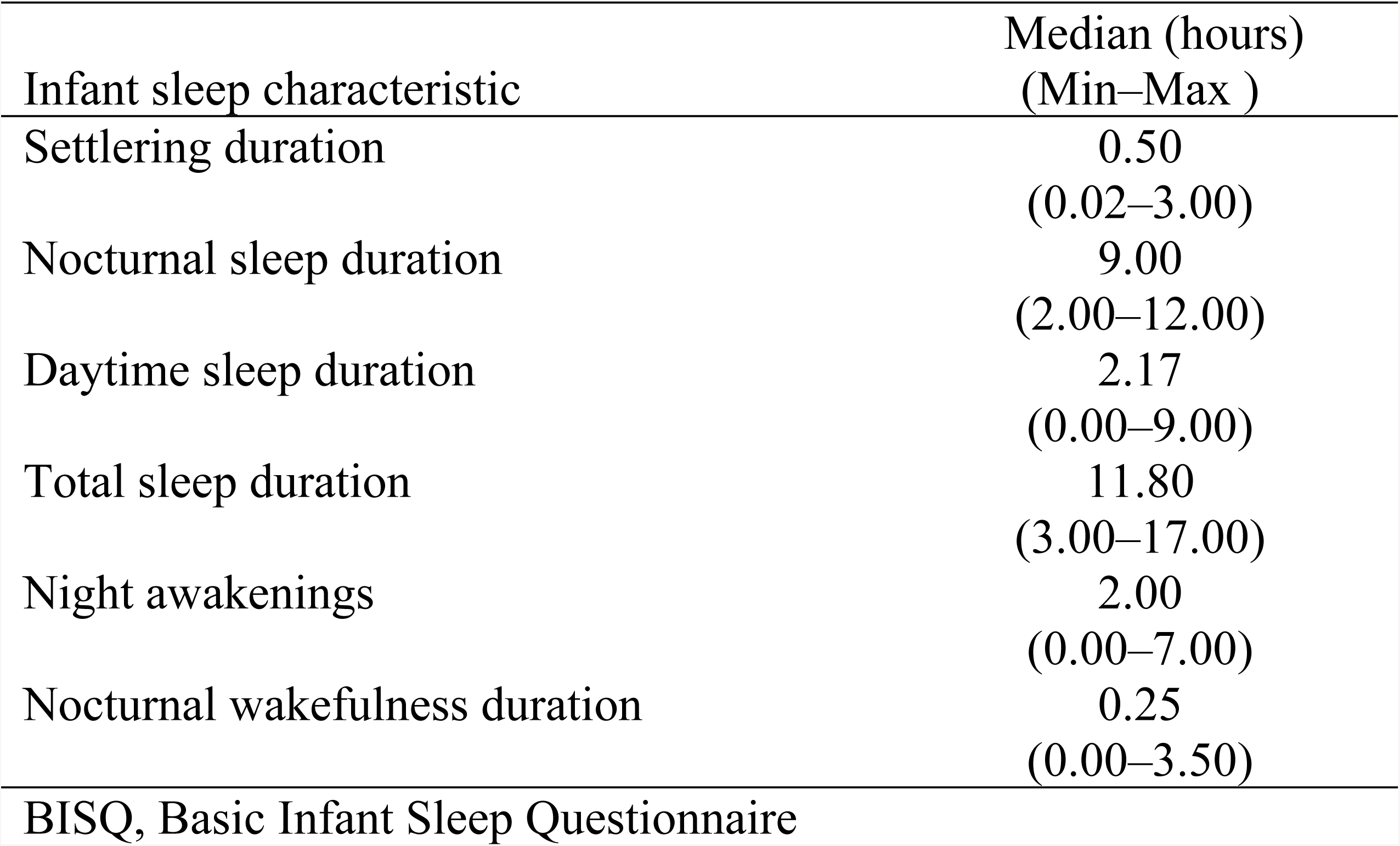
BISQ measurement results in general.

**Table 3.** Infant sleep characteristics (n=63)

| Sleep characteristic | Total |  |
| --- | --- | --- |
|  | n | % |
| Snoring |  |  |
| Never | 34 | 54.0 |
| Sometimes | 27 | 42.9 |
| Always | 1 | 1.6 |
| If the temperature is cold | 1 | 1.6 |
| Sleep location |  |  |
| Seperate room | 5 | 7.9 |
| Parent's room | 55 | 87.3 |
| Sibling's room | 1 | 1.6 |
| Other room in the house | 2 | 3.2 |
| Sleeping process |  |  |
| While feeding | 39 | 61.9 |
| In bed next to parent | 22 | 34.9 |
| Rubbed | 1 | 1.6 |
| Sleep position |  |  |
| supine | 10 | 15.9 |
| prone | 5 | 7.9 |
| side | 15 | 23.8 |
| combine | 33 | 52.4 |
| Cause of night awakenings |  |  |
| Hunger | 38 | 60.3 |
| bedwetting | 8 | 12.7 |
| Wants to play | 3 | 4.8 |
| Cold/hot temperature | 8 | 12.7 |
| Noises in bedroom | 1 | 1.6 |
| No answer | 5 | 7.9 |
| Toddler sleep quality |  |  |
| Normal | 42 | 66.7 |
| Problem | 21 | 33.3 |
| Mother's perception of her toddler's sleep quality |  |  |
| Excellent | 12 | 19.00 |
| Good | 29 | 46.00 |
| Fair | 17 | 27.00 |
| poor | 5 | 7.90 |
| Mother's perception of sleep problem in toddler |  |  |
| Very serious problem | 20 | 31.7 |
| Small problem | 16 | 25.4 |
| Not a problem at all | 27 | 42.9 |

In this study, factors that were significantly related to infant sleep quality included mother-infant bonding, infant stress level and parity; maternal depression and anxiety did not demonstrate any significant associations (Table 4). Therefore, only three factors were run in the logistic regression analysis to elaborate the magnitude of risk factors associated with problematic infant sleep quality. The goodness of fit test for the logistic regression analysis was good because the p-value of the Hosmer and Lemeshow test was 0.601, and the Omnibus test of model coefficients p-value was < 0.05. The odds ratios (ORs) between abnormality of mother-infant bonding and infant stress toward problematic infant sleep quality were 1.66 and 1.29, respectively (Table 5). Logistic regression analyses revealed that abnormality in the mother-infant bonding and infant stress predicted a 38.7% risk for problematic infant sleep quality (Nagelkerke R-squared, 0.387) (Figure 1). The number of parity was excluded in logistic regression analysis step 2 because it did not fulfil further criteria for logistic analysis (p>0.05 and included as variable not in the equation).

**Table 4.**
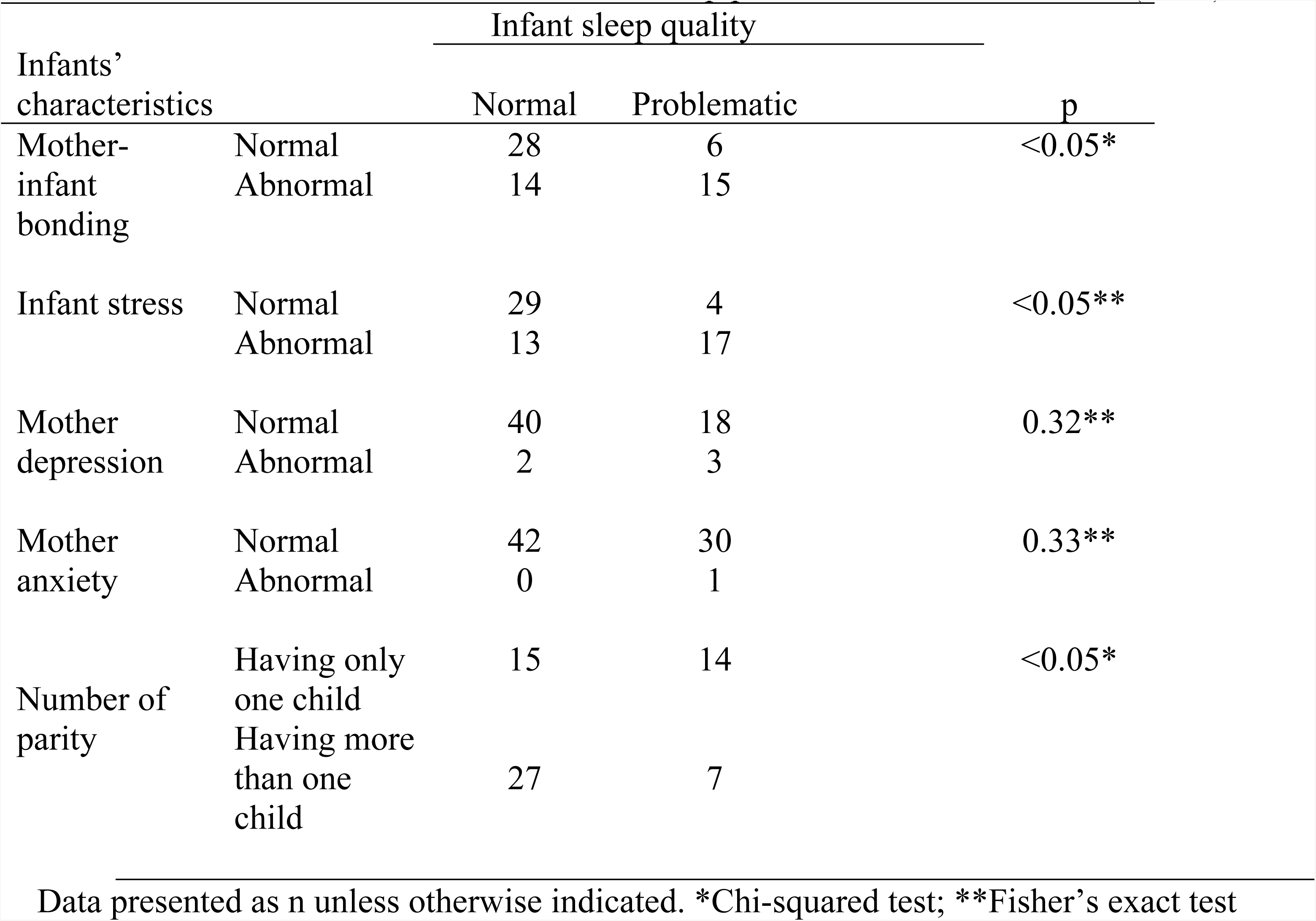
Bivariate association between infant sleep problems and related factors (n=63)

**Table 5.**
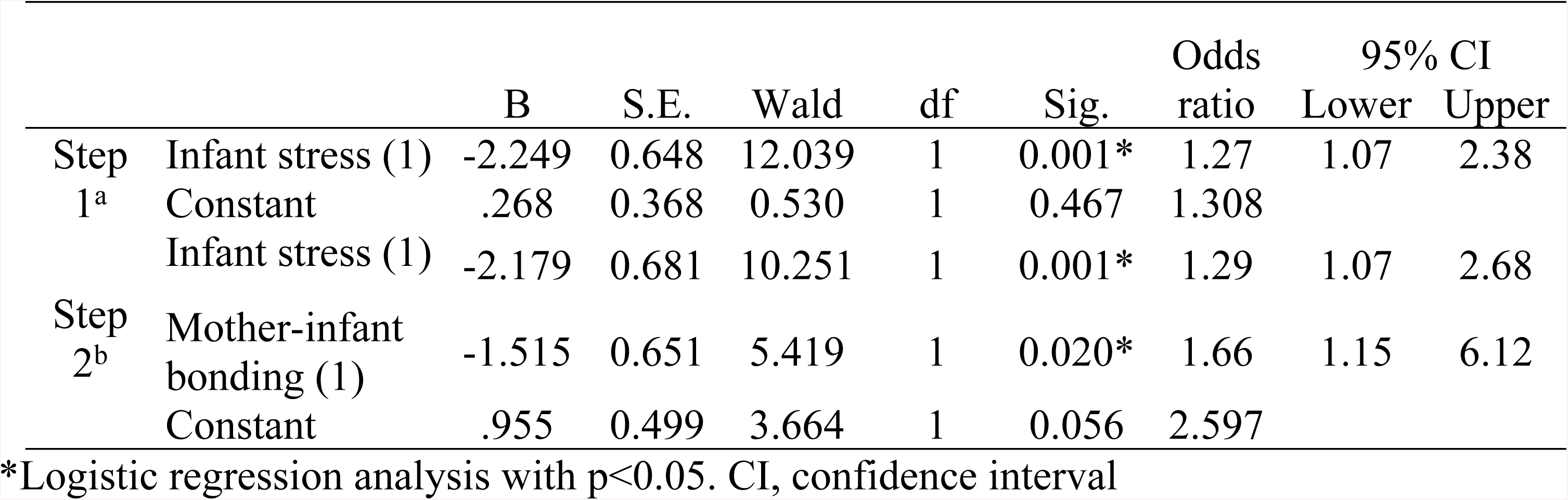
Association between abnormality in mother-infant bonding and infant stress toward problematic infant sleep quality (n=63)

**Fig 1.**
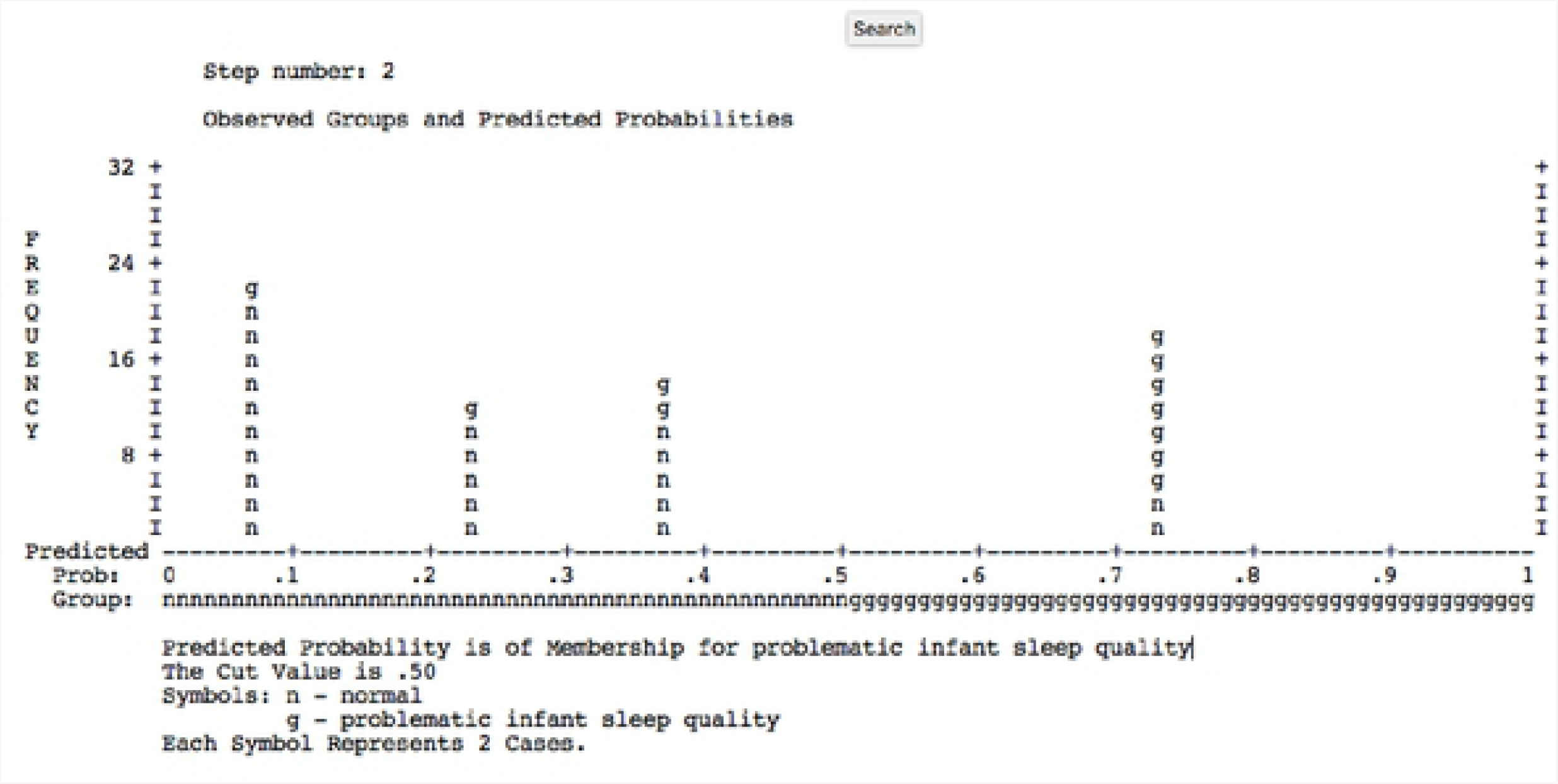
Observed group and predicted probabilities for problematic infant sleep quality.

## Discussion

Approximately 87.7% of infants sleep in the same room as their parents, and 63.1% fall asleep with the help of feeding. Ecology analysis of sleep in this study yielded a result similar to a previous study in which Indonesian toddlers were more likely to sleep either in a shared room or bed [12]. Asian parents also have a tendency to help toddlers initiate sleep [13]. Nocturnal and daytime sleep duration were 9 h and 2.17 h, respectively, with 2 awakenings and 0.25 h of nocturnal wakefulness. These results are similar to those from a previous study, which reported that Indonesian toddlers sleep 3.41 h during the daytime and 9.15 h at night, with 1.97 nocturnal awakenings and wakefulness for 0.68 h [12]. Based on the sleep disturbance criteria mentioned earlier, 33.3% of toddlers in this study had a sleep problem. This result was similar to a previous study reporting that 20–30% of toddlers worldwide and 33.3% of Indonesian toddlers experience sleep problems. The results indicated that there was a decrease in daytime sleep duration and the number of awakenings. As the toddlers grew, their sleep pattern started to concentrate at night [2]. Naturally, infants (7–18 months of age) experience fewer awakenings and start to exhibit a continuous sleep pattern [14]. Maturation of the circadian rhythm also plays a role. Newborns tend to sleep through the day because their circadian rhythm is not established. Starting at 2 or 3 months of age, the circadian rhythm starts its regulation and becomes more like the sleep pattern of adults [7].

Our results demonstrated that mother-infant bonding and infant stress level were two major risk factors that significantly affected infant sleep quality. Maternal instinct, infant intrinsic qualities, and mother-infant interaction factors have been known to contribute to infant sleep quality, and were predicted to have involvement in mother-infant bonding and the infant sleep pattern [4,15]. A previous theory—the transactional model—posits that sleep is mediated by the mother-infant interaction [15]. Disturbed interaction affects the mother’s parenting style and may cause sleep problems [16]. Parenting itself is difficult and challenging because infants do not generally exhibit efficient and comprehensive communication abilities. Thus, a mother needs to activate her limbic system to empathize and have a better understanding of her infant [14]. Bonding between mother and infant helps them to adjust to biopsychosocial adaption, which affects infant sleep pattern development [15]. A diminished mother’s responsiveness toward the infant is reflected in their emotional relationship and may also cause negative bonding [17].

Bonding is reflected though an affective behavior, such as soft voice, singing a lullaby or similar acts, which make infant feel comfortable, happy, and safe. A factor of the mother’s affective behavior is the maternal level of oxytocin (OT) and cortisol (CT). It has been found that mothers with high levels of OT and low levels of CT may be correlated with affective behaviors, such as motherly sound, positive maternal affect, and the touch of love [17-19]. Alternately, high maternal CT level is associated with a high infant CT level [20-22]. When an infant experiences non-affective acts, such as undesirable parenting or the lack of maternal affection, he/she may experience stress. Stress is the neurobiological response to adaptive function disruption, and may negatively impact a child’s development. It is related to increases in CT levels, which induces sympathetic responses in the body, such as elevated heart and respiratory rates, elevated blood glucose levels, and suppressed digestive function. In addition, stressful experiences also decrease neural plasticity, and the infant may exhibit hyper-reactive responses to stress [23]. In advance, infants tend to internalize events more easily, which makes it easier for them to develop depression or anxiety, which in turn leads to problematic sleep quality. It is also explained by the hypothalamus-pituitary-adrenal axis, which is activated by the stress response and CT is the output [23-25]. Acute and chronic stressors may cause changes in the CAR; for example, the awakened state from sleep is caused by a steep increase in the level of CT (38–70%) in the bloodstream. Although the function of this condition is not as clear, CAR has been predicted to play an important role in the awakened state by increasing place, time, and self-orientation [6].

The association between mother-infant bonding and infant sleep quality is complex. Although this study focused on the bonding impacts on infant sleep quality, other studies have focused on the opposite. In 2016, Hairston et al. [26] found that the problematic infant sleep quality at 5–8 months of age negatively affected mother-infant bonding. Early sleep consolidation can facilitate better sleep patterns in the infant, which in turn are associated with better mother-infant bonding, parent well-being, family harmony, and infant temperament. Failure in sleep consolidation increases parent uneasiness and may change the interaction pattern to one of negative engagement [14,27]. The number of parities may contribute to the toddler’s sleep quality through the mother’s experience and knowledge of caretaking. A few studies have shown that multiparous mothers demonstrate a better parenting characteristics than primiparous mothers [9]. Parents’ lack of knowledge of the infant’s sleep needs causes them to experience difficulties with identifying possible infant sleep problems [28]. Moreover, the parents’ unawareness of the toddler’s sleep patterns may cause a lower level of communication with a health provider and a higher sleep problem risk for the toddler [29]. Primiparous parents have been found to be more responsive toward a child crying and are capable of soothing effectively [30]. High parent intervention during the infant sleep process is associated with a higher number of night-time awakenings [31]. External intervention reduces an infant’s chances at proper sleep self-regulation [4]. The parent’s self-limiting of the infants’ cries is required to let the toddler learn how to sleep by themselves [32]. Alternately, a mother ought to know their child’s signs to provide appropriate emotional responses or emotional availability (EA). High maternal EA during sleep time has been predicted to increase the number of better sleep time quantity. Maternal EA helps toddlers feel safe and trust their environment [33]. The mother’s sensitivity level in understanding the toddler plays a role in the toddler’s sleep. In a study, mothers who worried that their children were experiencing sleep problems demonstrated more accuracy in identifying the infant sleep problems than those who believed their children slept normally. This illustrates that maternal insensitivity can become a risk factor in infant sleep problems [4]. However, in our study, the mothers’ depressive and anxious condition was not associated with problematic infant sleep quality. This may be explained that our samples mostly lived in extended families and developed an optimal bonding quality with the infant; however, future studies are warranted.

In conclusion, the prevalence of sleep problems in toddlers was high (33.3%), and need more attention due to the high percentage of parents who did not consider sleep problems to be particularly concerning (42.9%). Results of this study implied an association between infant sleep quality problems and mother-infant bonding and infant stress. Nevertheless, this study had several limitations, such as the lack of cut-off of MIBS score in infant and infant stress level; therefore, this study applied the Z-score to categorize the abnormal state of those conditions. Furthermore, this study did not include several other factors, such as family harmony, infant temperament, CT level, among others, which may be associated with infant sleep quality. However, this study was the first in Indonesia to elaborate mother-infant bonding and infant stress level in infant sleep quality. Therefore, it can be used as basic data for further research in the field of infant mental health. In addition, the results of the present study may also be used to improve infant mental health services by disseminating the mother-infant bonding scale and infant stress questionnaire in primary health centers, especially to detect problems and manage them earlier because brain neuroplasticity among infants is still in development. Thus, early screening of mother-infant bonding and infant stress level questionnaires are required in community health services, such as integrated service post (Posyandu) in Indonesia, to achieve better sleep and outcomes of child’s growth and development.

## Acknowledgements

This study was supported by the Faculty of Medicine Universitas Indonesia Research Grant 2018 (Hibah PITTA 2018). The authors thank all the mothers and infants who participated in this study.

